# Hock immunization: a refined alternative for developing Experimental Autoimmune Myasthenia Gravis (EAMG) in mice

**DOI:** 10.1101/2025.03.06.640628

**Authors:** Marina Mané-Damas, Anja K Schöttler, Tanya Mohile, Britt Arets, Erdem Tüzün, Mario Losen, Pilar Martinez Martinez

**Author notes:** Corresponding author: Pilar Martinez Martinez, Department of Psychiatry and Neuropsychology, Mental Health and Neuroscience Research Institute, Faculty of Health, Medicine and Life Sciences, Maastricht University, Maastricht, the Netherlands.

## Abstract

Experimental autoimmune myasthenia gravis (EAMG) is an active-immunization model that was first discovered by coincidence when rabbits injected with purified acetylcholine receptor for neuromuscular junction physiology studies, suddenly developed muscle weakness. In mice, this model typically involves bilateral subcutaneous injections of purified torpedo acetylcholine receptor (tAChR) in the hind footpads and the scapular regions, followed by one or two booster immunizations in the thighs and over the scapulas. However, footpad injections can cause severe discomfort and progressive debilitation. Hock immunization has shown to be a viable alternative in other models, offering comparable efficacy without impairing mobility caused by inflammation, thus reducing discomfort.

To investigate the use of hock immunization in the mouse EAMG model, 7-week-old female C57BL6/6J (B6) mice were bilaterally injected over the scapulas and either in the hocks or footpads for the primary immunization. All animals received a booster immunization 4 weeks later, following standard guidelines.

Inverted mesh measurements demonstrated that tAChR-immunized animals whether receiving hock or footpad injections, exhibited similar muscle weakness. This weakness correlated with reduced body weight, which reached statistical significance, only in the hock-immunized group. tAChR antibodies were detectable in both groups 4 weeks after primary immunization and continue raising until the end of the experiment. Additionally, tAChR immunized mice required significantly less curare to achieve decrement in muscle response. Notably, hock-immunized mice displayed consistent mechanical sensitivity over time, which was significantly lower compared to footpad-immunized mice.

In conclusion, hock-immunization is a refined and effective alternative for developing the EAMG mouse model. It results in similar disease incidence and severity while minimizing immunization-related discomfort compared to the standard footpad method.

## Introduction

Myasthenia gravis (MG) is a classic antibody-mediated autoimmune disorder, caused by autoantibodies targeting the acetylcholine receptor (AChR) in most cases, or other molecules involved in clustering the receptor at the neuromuscular junction (NMJ) ^1^. These autoantibodies reduce the number of AChRs available at the membrane through various mechanisms, including antigenic modulation, receptor internalization, and complement activation, ultimately, resulting in impaired neuromuscular transmission. The extensive knowledge available on MG pathophysiology makes it a valuable disease model for investigating novel therapeutic approaches, not only for MG, but also for other antibody-mediated autoimmune disorders with shared mechanisms ^2^.

Experimental autoimmune myasthenia gravis (EAMG) is an active-immunization model discovered when rabbits injected with purified *Electrophorus electricus* AChR ^3-5^ to raise antibodies for NMJ physiology studies, unexpectedly developed muscle weakness ^4^. EAMG closely mimics the clinical, immunopathological and electrophysiological features of human AChR-MG, making it an attractive model for drug testing and further exploration of MG pathophysiology.

In mice, the model typically requires bilateral subcutaneous injection of *Torpedo californica* AChR (tAChR) emulsified with Complete Freund’s Adjuvant (CFA) into the hind footpads and scapular regions, followed by one or two booster immunizations in the thighs and scapular regions 4 and 8 weeks later ^6^. The footpads are considered desirable immunization sites because the lymph drainage leads directly to the popliteal, medial iliac and inguinal lymph nodes ^7-9^. However, because footpads are weight-bearing structures, the inflammation and swelling solely caused by immunization can lead to unrelieved stress, potential lameness, severe discomfort, and progressive debilitation. In line with the EU directive 2010/63/EU, which emphasizes refinement of animal models, we propose using the hocks as an alternative immunization site for the EAMG mouse model. The hocks, located in the lateral tarsal region just above the ankle, utilize the same lymph nodes as the footpad while avoiding interference with mobility and reducing discomfort ^10^. To the best of our knowledge, this immunization method is not included in standard guidelines and has never been described for MG.

These observations encouraged us to investigate the efficacy of hock immunizations in the EAMG mouse model and compare it to the standardized model using footpad immunizations. Since the main difference between the two models is the primary immunization site, the newly proposed alternative model, will be called **hock-20 or 40 µg tAChR** (EAMG) and the standardized model will be referred to as **footpad-20 µg tAChR** (EAMG) from now on. Animals injected with saline will be referred to as **hock-** or **footpad -saline** (nonMG).

We first tested 20 and 40 µg tAChR doses in the hock regions and compared it to the standard dose of 20 µg tAChR in the footpad to select the best EAMG-inducing dose for the hock-immunization animals (Table 1). Once the optimal tAChR dose for hock immunization was identified, we compared the disease incidence, severity, and discomfort for hock and footpad-EAMG, to saline-injected hock- and footpad, nonMG control animals (Phase 1 experiment, Table 2). In this second study, some methods and parameters used to assess muscle weakness were adapted to increase sensitivity.

**Table 1.** Animal groups and doses phase 1 experiment.

| Group | Hock/footpad saline/tAChR | Immunization mix | Primary Immunization site (week 1) | Booster Immunization site (week 5) | Number of Animals |
| --- | --- | --- | --- | --- | --- |
| 1 | Hock-saline (nonMG) | Saline: CFA | Hind hocks and in scapular region | Thighs and in scapular region | 12 |
| 2 | Footpad-saline (nonMG) | Saline: CFA | Hind footpads and in scapular region | Thighs and in scapular region | 12 |
| 3 | Hock-20 µg tAChR (EAMG) | 20 µg tAChR: CFA | Hind hocks and in scapular region | Thighs and in scapular region | 12* |
| 4 | Hock-40 µg tAChR (EAMG) | 40 µg tAChR: CFA | Hind hocks and in scapular region | Thighs and in scapular region | 12 |
| 5 | Footpad-20 µg tAChR (EAMG) | 20 µg tAChR: CFA | Hind footpads and in scapular region | Thighs and in scapular region | 12 |

\*This includes an animal that was euthanized due to severe dermatitis in the scapulas and has not been considered for further analysis.

**Table 2.** Animal groups and doses phase 2 experiment.

| Group | Hock/footpad saline/tAChR | Immunization mix | Primary Immunization site (week 1) | Booster Immunization site (week 5) | Number of Animals |
| --- | --- | --- | --- | --- | --- |
| 6 | Hock-saline (nonMG) | Saline: CFA | Hind hocks and in scapular region | Thighs and in scapular region | 10 |
| 7 | Footpad-saline (nonMG) | Saline: CFA | Hind footpads and in scapular region | Thighs and in scapular region | 10 |
| 8 | Hock-20 µg tAChR (EAMG) | 20 µg tAChR: CFA | Hind hocks and in scapular region | Thighs and in scapular region | 20 |
| 9 | Footpad-20 µg tAChR (EAMG) | 20 µg tAChR: CFA | Hind footpads and in scapular region | Thighs and in scapular region | 20 |

## Results

### Survival and clinical manifestations are similar between hock- and footpad-immunized animals with the same tAChR dose

Animals injected with 20 µg tAChR, regardless of the immunization site, showed a ∼10% drop-out due to severe clinical manifestations which led to earlier termination or HEP (Figure 2a) as previously observed in other EAMG models ^11^. Immunization with 40 µg tAChR resulted in lower survival of 75%. One animal from the hock-20 µg tAChR group in phase 1 experiment developed severe dermatitis over the scapulas and was removed from the experiment. This animal has not been considered for further analysis, as the inflammation could interfere with the immune response and therefore the development of the EAMG model.

**Figure 1.**
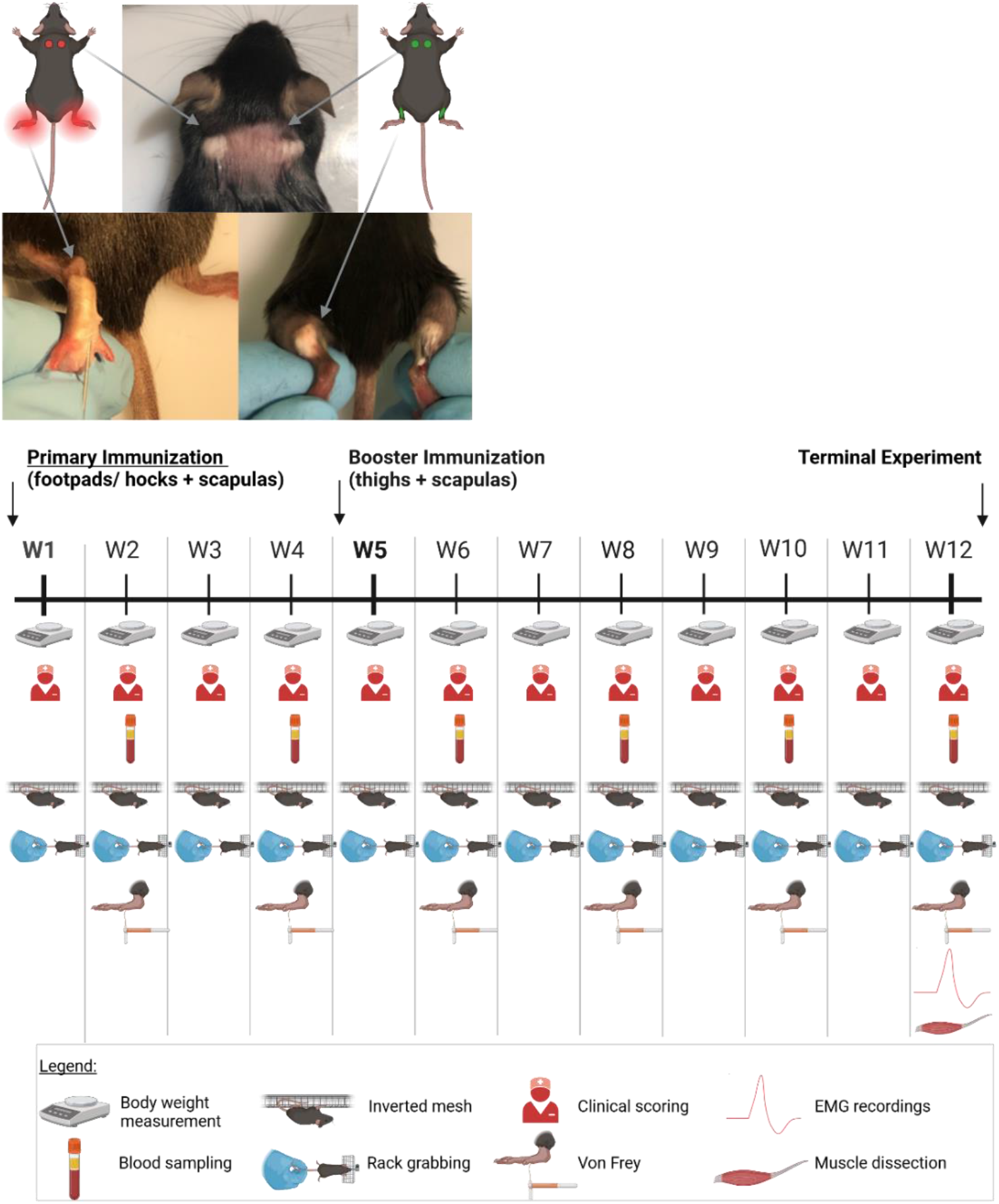
Alternative immunization strategy for the EAMG mouse model: study rationale and timeline. Animals were bilaterally immunized in the footpads or hocks and over the scapulas at the beginning of week 1, with booster immunizations in the thighs and over the scapulas at week 5. All animals underwent regular tests, including body weight measurement (weekly), clinical scoring (2 times per week) and muscle strength tasks (inverted mesh and rack grabbing, weekly). Additionally, blood sampling and von Frey measurements were conducted every other week. Before euthanasia (typically at the end of week 12), animals underwent electromyography with curare infusion (if required), and muscle dissection for post-mortem analyses. Created in https://BioRender.com

**Figure 2.**
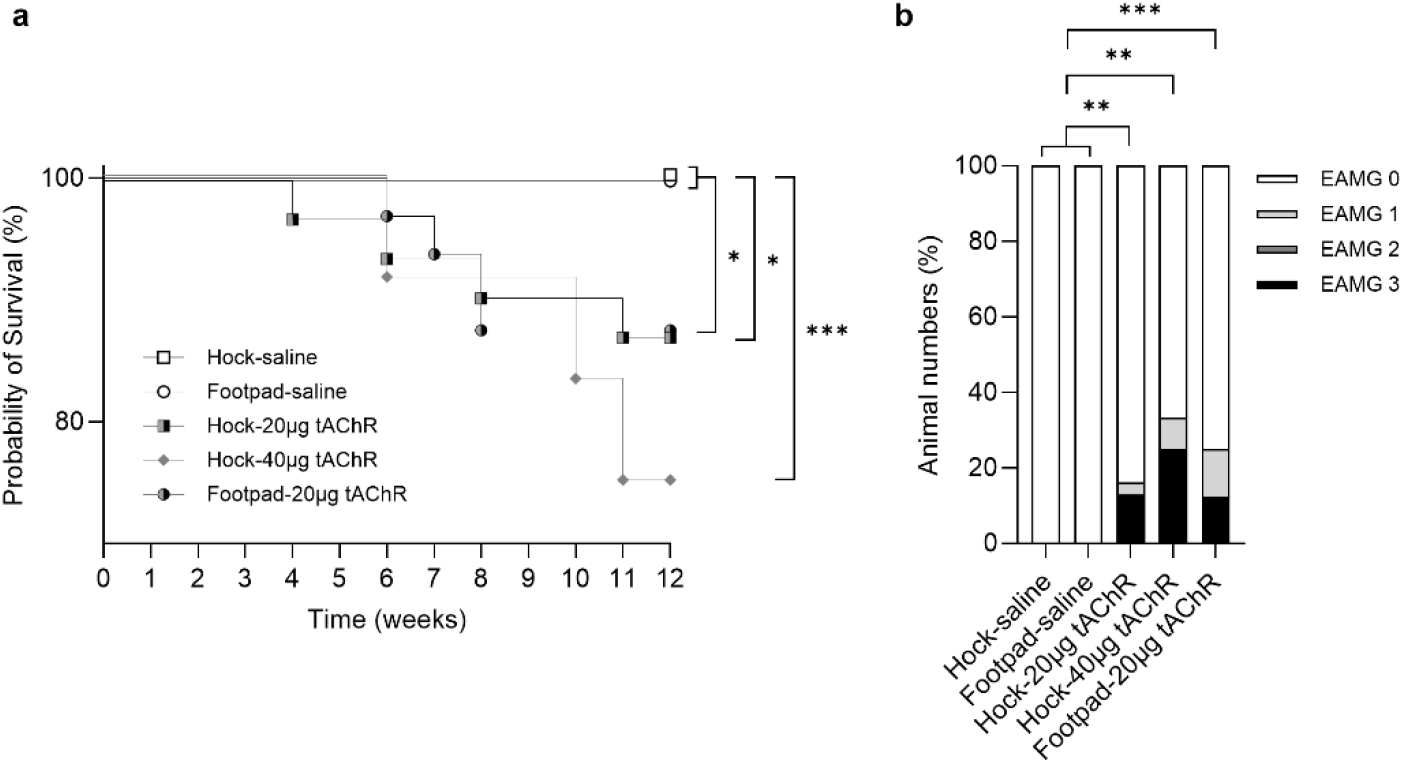
Survival analysis and EAMG clinical scores. Animals from phase 1 and phase 2 experiments were grouped (see Table 1 and 2) for **a)** survival data, where, as expected, the hock-40 µg tAChR group had the highest drop-out rate compared to the lower dosed groups of 20 µg tAChR, both in the footpads and the hocks. All tAChR-injected animals had lower survival probabilities compared to saline-injected animals. Hock and footpad saline-injected animals were combined for this analysis. Log-rank (Mantel-Cox) test for all groups [p=0.0332] and Log-rank (Mantel-Cox) test for individual comparisons, with statistically significant differences indicated in the graph. **b)** Clinical scores of muscle weakness were assessed by a blinded investigator at the terminal timepoint [12 weeks after primary immunization or earlier if animals reached HEP]. 0 = no abnormalities, 1 = fatigable weakness, 2 = constant weakness, 3 = severe muscle weakness or more than 20 % body weight loss. Fisher’s exact test for all [p= 0.0014] and for individual comparisons, with statistically significant differences indicated in the graph. *p<0.05, **p<0.01 and ***p <0.001.

Regarding clinical manifestations, tAChR immunized animals had significantly higher EAMG scores compared to saline immunized animals (Figure 2b). The proportion of animals with severe clinical manifestations (EAMG score of “3”) was higher in the 40 µg tAChR immunized animals compared to the 20 µg tAChR immunized animals, both in the hock and footpad groups, as expected.

### Hock immunization with tAChR induces significant body weight loss

Body weight was not a strong readout parameter for the mouse EAMG model, as animals continued to gradually gain body weight during the experiment (Figure 3a). Significant body weight differences were only observed before euthanasia, where animals injected with 20 µg tAChR in the hocks experienced significant body weight loss compared to saline-injected animals (Figure 3b).

**Figure 3.**
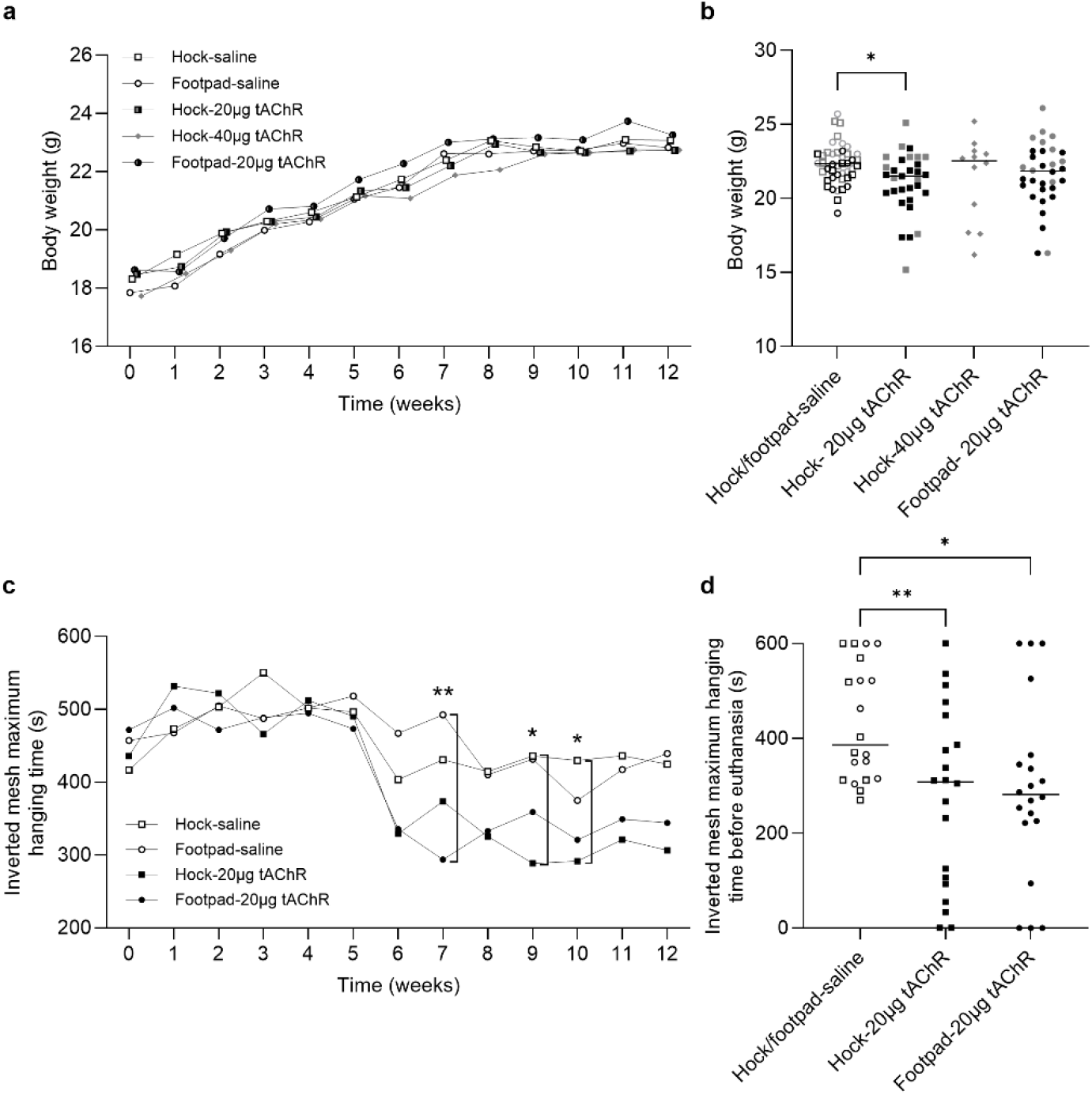
Mouse body weight and muscle strength. Mouse body weights were grouped together for all animals tested in both experiments (see Tables 1 and 2). **a)** Weekly body weight data of all groups for 12 consecutive weeks (average/group) and **b)** individual body weights at the terminal timepoint [12 weeks after primary immunization or earlier if animals reached HEP]. Open and filled symbols represent saline and tAChR injected groups respectively. Grey and black symbols correspond to mice in the first and second phase studies respectively. **c)** Weekly maximum time spent by the animals in the inverted mesh (data exclusively from the phase 2 experiment with hanging time adapted to 600 s) for 12 consecutive weeks (average/group) and d) individual hanging times at the terminal timepoint [12 weeks after primary immunization or earlier if animals reached HEP]. Hock and footpad saline-injected animals were combined for this analysis. One and two-way ANOVA, with multiple comparison of the indicated groups and Bonferroni post hoc testing were used for statistical analyses. *p<0.05 and **p<0.01.

### Hock- and footpad tAChR immunized animals showed significantly increased muscle weakness

After booster immunization (week 5), both hock and footpad tAChR immunized groups showed significantly reduced muscle strength which persisted until the end of the experiment (Figure 3c, phase 2 experiment, 600 s maximum hanging time). Significant differences in maximum hanging times were observed at week 7 between the footpad-20 µg tAChR and footpad-saline groups and at weeks 9 and 10 between hock-20 µg tAChR and hock-saline groups (p<0.05, One-way ANOVA and Bonferroni post hoc test). At the terminal timepoint (12 weeks after primary immunization or earlier if animals reached HEP), both hock and footpad tAChR immunized groups showed significantly reduced maximum hanging times compared to saline immunized animals (Figure 3d). Using 320 s as maximum hanging time, in the phase 1 experiment was less sensitive and therefore insufficient to show significant differences between the groups at any timepoint (Supplementary Figure 1a), except at the terminal timepoint whereas a significant difference was observed between animals injected with hock-40 µg tAChR and saline immunized animals (Supplementary Figure 1b). Average grip strength was also measured in the phase 2 experiment, but differences over time were subtle and variable (Supplementary Figure 2a). Significant differences were observed between hock-saline immunized animals and hock-20 µg tAChR immunized animals in week 9 (p<0.05, One-way ANOVA and Bonferroni post hoc testing) (Supplementary Figure 2b). Before euthanasia, saline immunized animals were significantly stronger than those immunized with tAChR regardless of the immunization site (Supplementary Figure 2c).

### tAChR autoantibody levels are detectable before booster immunization

Antibodies against tAChR are proven to be elevated in rodent EAMG models a factor 100 compared to rodent AChR autoantibody levels ^12^. tAChR antibody levels were slightly increased but already significantly higher 4 weeks post-primary immunization (Figure 4a) and continued rising after the booster immunization (Figure 4b) and until the end of the experiment, before euthanasia (Figure 4c) with no significant differences between hock- and footpad-tAChR immunized animals at any timepoint.

**Figure 4.**
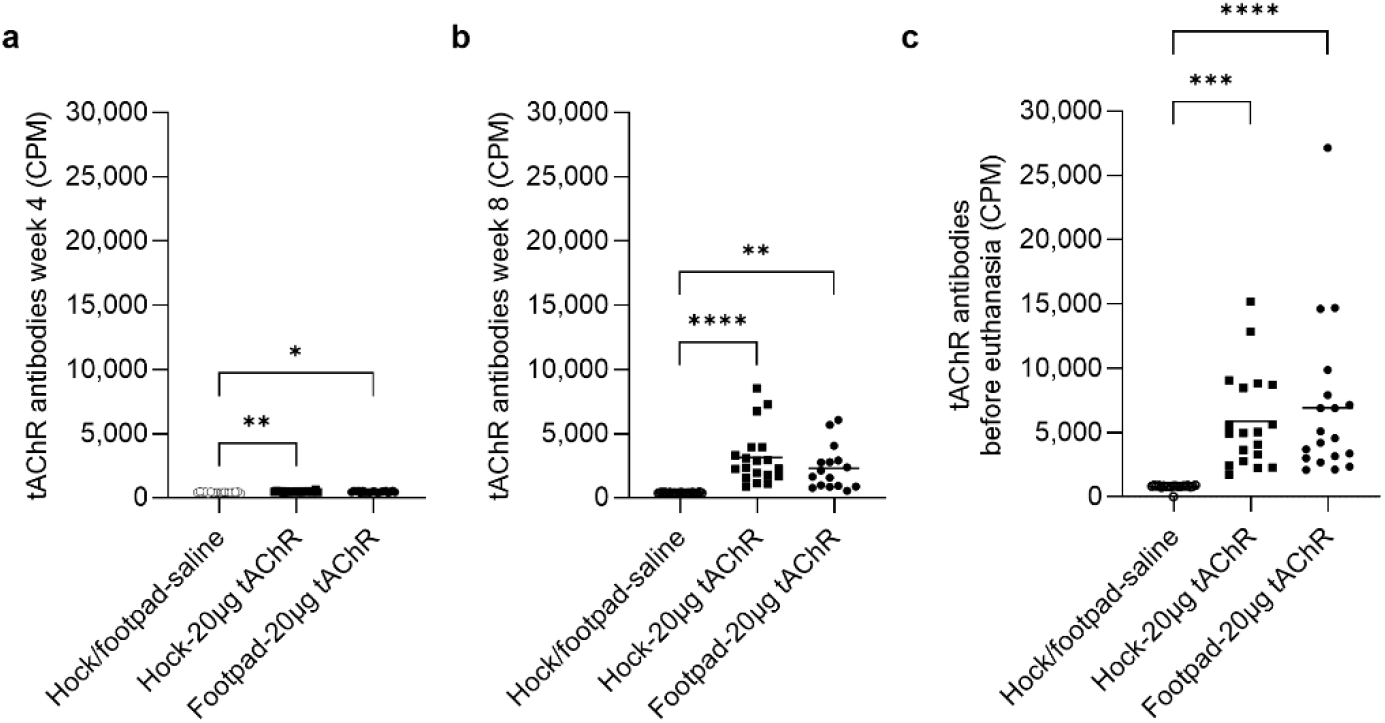
Plasma tAChR autoantibody levels. tAChR antibody levels at **a)** week 4, **b)** week 8 and **c)** terminal timepoint [12 weeks after primary immunization or earlier if animals reached HEP] from phase 2 experiment. One-way ANOVA, multiple comparisons of the indicated columns and Bonferroni post hoc testing were used for statistical analyses. *p<0.05, **p<0.01 and ****p<0.0001. CPM; counts per minute.

### tAChR immunized animals showed greater functional muscle AChR content loss

To detect the level of remaining functional AChR at the NMJ, we challenged the muscles with continuous IP infusion of curare, a toxin that competitively binds to available AChR and thereby inducing paralysis. tAChR immunized animals required significantly lower/no curare dose infusions to reach decrement, indicating a strong AChR loss in the muscles compared to saline immunized animals (Figure 5).

**Figure 5.**
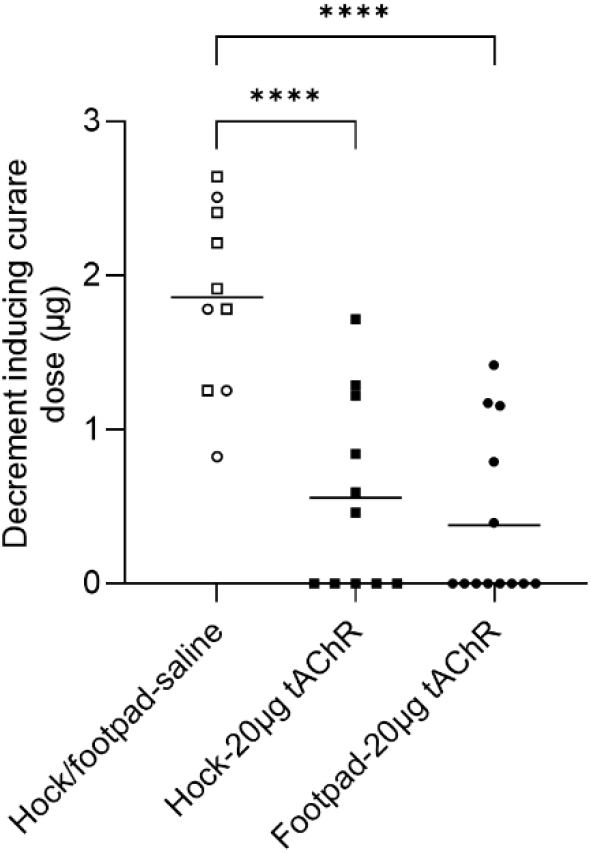
Muscle AChR content at terminal. Decrement-inducing curare dose during EMG as an indirect measurement of the amount of AChR in the muscles from phase 2 experiment. Only animals with 3 consecutive decrement measurements have been included. One-way ANOVA, multiple comparisons and Bonferroni post hoc testing were used for statistical analyses. ****p<0.0001.

### Hock immunization resulted in less discomfort compared to footpad immunization

With similar baseline levels to mechanical sensitivity, footpad-immunized animals soon showed increased sensitivity, which gradually decreased over time, while the sensitivity of hock-immunized animals remained relatively stable during the experiment (Figure 6a). Differences became more pronounced after immunizations in weeks 2 and 6. At week 2 post immunization, footpad-immunized animals exhibited significantly higher sensitivity to mechanical stimulus (Figure 6b), which was expected since sensitivity was tested in the footpads. However, these differences persisted over time. By week 6, footpad-immunized animals displayed significantly higher sensitivity to mechanical stimulus. In this case, the increased sensitivity cannot be attributed to the immunization site, as all animals received the booster immunizations at the same sites: scapulas and thighs (Figure 6c). tAChR and saline-immunized animals were pooled and grouped by immunization site.

**Figure 6.**
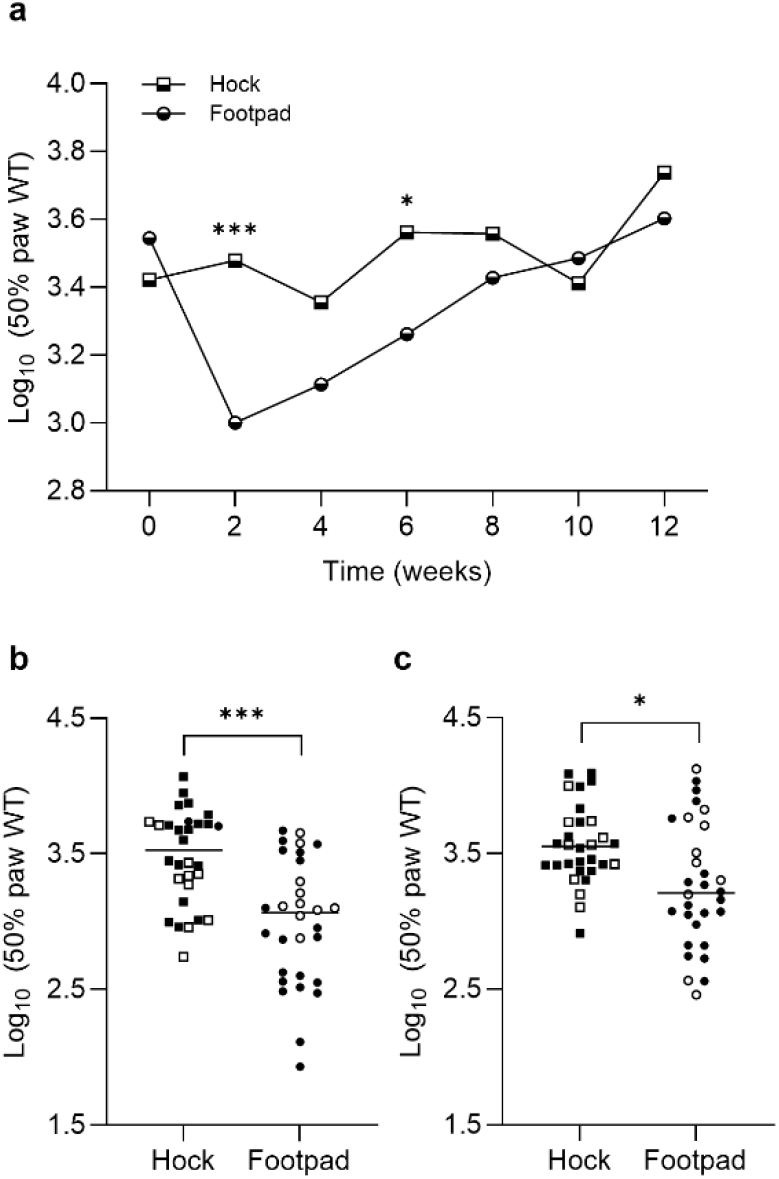
Injection site effect on mechanical sensitivity. Animals from the phase 2 experiment were grouped based on their immunization site: hock or footpad, regardless of the substance injected (saline or tAChR). Group-averaged Log_10_ (10,000 × 50% paw WT) values were **a)** longitudinally analyzed every other week for a total of 12 weeks, and individual values are shown for **b)** week 2 and **c)** week 6 post-immunization. One and two-way ANOVA, multiple comparisons and Bonferroni post hoc testing were used for statistical analysis. *p<0.05; ***p<0.001.

## Discussion

Animal studies in antibody-mediated autoimmune disorders, such as MG, have clarified disease mechanisms, leading to new therapeutic targets and improved treatments. The detailed study of AChR-MG also serves as a valuable model for similar disorders, providing proof of principle for broader research.

However, it is essential to ensure that animal use is restricted to research questions that cannot be answered without living organisms, using the minimum number of animals and applying refining strategies, to improve animal welfare whenever possible. In line with this, the EAMG rat model is a robust and widely accepted model involving a single administration of antigen, emulsified with CFA, effectively replicating many of the characteristic pathogenic features of MG ^12^. In contrast, the standardized EAMG mouse model requires a more discomfort-inducing procedure, involving multiple CFA injections, with the first administered in the footpads, a highly sensitive area ^6^. Given the advantages of mouse models, including the availability of genetically modified mouse lines, it is important to refine the EAMG mouse model to make it more humane.

In this study, we demonstrate that hock immunization, without a second booster, is a suitable strategy for developing the EAMG mouse model with reduced mechanical sensitivity compared to the standardized model. Using the same tAChR antigen and CFA amounts, both hock and footpad-immunized animals showed comparable antibody levels and functional AChR loss, with similar muscle weakness and fatigue. However, hock-immunized animals showed reduced sensitivity to mechanical stimuli overtime compared to footpad-immunized animals, suggesting reduced discomfort in the hock-immunized group.

Several researchers have explored alternative strategies for developing the EAMG mouse model. For example, different adjuvants, including incomplete Freund’s adjuvant (IFA), aluminum hydroxide, lipopolysaccharide (LPS) and Polyinosinic-polycytidylic acid (Poly(I:C)). However, the first two were insufficient for EAMG development ^13-17^. Mice immunized with LPS:IFA and tAChR in the footpads and scapulas, followed by two booster immunizations, showed clinical manifestations similar to the standardized model ^18^. The authors found that the LPS response was CD4-independent unlike CFA with heat-killed *Mycobacterium tuberculosis*, which requires CD4 cells for an immune response. The isotypes developed were mainly of the IgG2b and IgM in the LPS induced model, while IgG2b and IgG1 but not IgM contributed to the pathology in the animals immunized with CFA. However, this approach did not aim to refine the model to reduce discomfort. It could be worthwhile to explore the effect of LPS in the newly developed hock-EAMG mouse model to see differences in the disease induction and severity. In the case of poly(I:C) animals injected intraperitoneally every 3 days for 6 weeks, with the adjuvant, developed a milder EAMG phenotype with muscle weakness and trace amounts of AChR autoantibodies ^19^. However, frequent intraperitoneal injections cause severe discomfort, and may lead to peritonitis, potentially altering immune response and interfering with MG model readouts.

Other approaches have tested different antigen sources. *Escherichia coli* plasmids expressing recombinant human AChR alpha or gamma subunits efficiently induced ocular symptoms in rodents but did not produce generalized MG ^20-22^. More recently, Theissen and colleagues used recombinant AChR to develop EAMG in mice. Immunization with 40 μg of the AChR alpha subunit (CHRNA1) emulsified with CFA in the flanks and tail base, followed by a booster led to the presence of AChR autoantibodies, complement deposition at the NMJ and clinical signs of muscle weakness and fatigue measured by grip strength ^23^. Like our model, clinical manifestations developed 2 weeks post-booster immunization. While avoiding footpad injections may reduce discomfort, using only the alpha subunit limits translational relevance. About half of the antibodies detected in AChR-MG patients target the alpha subunit. However, autoantibodies against the other four subunits of the receptor (including the gamma subunit of the fetal AChR) have been found, even in the same patients with anti-alpha AChR antibodies ^24-27^, with variable percentages in the abundance of antibodies against each of these subunits. Despite an animal study showing a stronger pathogenicity by anti-alpha AChR antibodies in rats compared to those against other subunits ^28^, the relevance of these antibodies into the pathogenesis of AChR-MG in humans is not yet clear. Using the full AChR molecule such as from *Torpedo californica* or human extracellular domains of various subunits as proposed by Lazaridis and colleagues, offers a more physiological response ^29^.

During the development of the model, we noted several key observations for assessing the EAMG mouse model. In our study, the 600 second inverted mesh test with pre-exercise was the most sensitive method for measuring muscle weakness. Contrary to other studies, grip strength lacked sensitivity, showing few differences between disease and non-diseased animals. Additionally, unlike in rat models, body weight loss was not a reliable indicator of MG disease incidence or severity. We also highlight the importance of EMG with curare infusion as a robust, sensitive method for evaluating disease severity in rat and mouse EAMG models. Intermediate EMG measurements without curare infusion could also serve as additional readout to assess functional AChR loss over time.

A limitation of our study is the assessment of discomfort using Von Frey testing which involves the footpads. However, the use of the von Frey is a direct measure of the discomfort that the animal is experiencing while performing regular activities such as walking, with a good translational value of the overall discomfort of the animal. Other pain assessment methods, such as the Hargraves test, also use footpads to measure thermal sensitivity or could be influenced by the muscle weakness developed in our model. As expected, mechanical sensitivity in footpad-immunized animals was significantly higher after immunization, and remained elevated until week 10 post-immunization, at which point it aligned with hock-immunized animals. In contrast, hock immunization did not alter mechanical sensitivity, indicating no significant discomfort or pain in these animals.

In conclusion, hock immunization is a suitable and more humane alternative to footpad immunization for developing the EAMG mouse model. This approach reduces discomfort while maintaining the disease characteristics necessary for effective research.

## Methods

### EAMG induction

Female C57BL6/6J (B6) mice were purchased from Charles River. The experiments were approved by the Committee on Animal Welfare (Project License number 2016-005) in accordance with European and Dutch laws, rules, and guidelines (86/609/EU).

Seven-week-old mice were housed in individually ventilated cages in groups of 3. EAMG was induced with tAChR extracted and purified from *Torpedo californica* as previously described ^30^. Briefly, 20 or 40 µg tAChR were emulsified with an equal volume of CFA containing 0.1% of *Mycobacterium tuberculosis* to a final volume of 200 µL. Primary immunization used bilateral injections in the hind footpads or in the hocks, and over the scapulas (50 µL per immunization site). Animals received a booster immunization 4 weeks later, using the same immunization mix and volume, this time in the thighs and over the scapulas (more distal compared to the first immunization to prevent inflammation/irritation). Control animals (nonMG) received a 1:1 mix of saline and CFA during immunizations, following the same injection sites and volumes as the EAMG animals. To ensure consistency in immunogenic capacity, all animals received the same tAChR mix and CFA from the same batch (Table 1 and 2). Animals received 1 mg/kg buprenorphine orally via drinking water for a week after the primary and booster immunizations. All animals were euthanized 12 weeks after the primary immunization unless a humane endpoint (HEP) was reached earlier.

### General body condition and clinical scoring

Animals were observed daily and weighed weekly, or more often if required, to assess their general condition and record potential muscle mass loss related to the disease model. Animals with body weight loss >10% were provided with gel booster food, wet pellets and long nipples. Subcutaneous injections of saline were also administered if signs of dehydration were observed.

Clinical scoring was conducted twice weekly by two independent researchers as described previously ^6^. Briefly, mice were placed on the roof of the cage and pulled backwards along the bars by the base of the tail, ensuring that all four paws were gripping the cage as it is the natural reflex. After twenty times repetition, they were observed for any signs of muscle weakness and fatigue. Clinical scoring was graded as follows: “0”= no abnormalities observed; “1”= weakness evident only after exercise; “2”= visible clinical signs before/during exercise; “3”= severe weakness, dehydration, paralysis (quadriplegic) and loss of significant body weight (> 15% maximum body weight); “4”= found dead in the cage. The HEP was reached when an animal lost ≥20 % of its maximum body weight and had a clinical score of 3; any such animal was euthanized within 24h.

### Motor function tests

Animals were weekly tested for muscle weakness and fatigue as an index of disease incidence and severity, as described previously ^6^. Briefly, all animals were tested for their ability to; 1) repeatedly grasp a metal grid attached to a grip strength meter (GRIP GS3 PanLab Harvard medical) using their front paws, while held manually near the base of the tail and retracted manually (rack grabbing) for 5 consecutive times and 2) remain upside down in a mesh, placed ∼50 cm above the ground, for a maximum of 320 s in phase 1 and 600 s in phase 2 experiment (inverted mesh). To increase sensitivity, all animals were pre-exercised before these tasks in the phase 2 experiment as previously described ^6^.

### Analysis of mechanical (pain) sensitivity

Mechanical sensitivity was assessed by plantar application of calibrated Von Frey filaments to the left and right hind paws, logarithmically increasing in force (0.008, 0.04, 0.07, 0.16, 0.4, 0.6, 1.0, 1.4, 2.0, and 4.0 g) using a modified version of the up-down method described by Chaplan [49]. Only withdrawal responses associated with sharp withdrawal or aversive behavior to the stimulus (e.g. paw licking, postural changes, and/or attacking the filament during paw stimulation) were considered a positive response. After a positive withdrawal reaction, two additional positive withdrawal reactions were recorded. The 50% paw withdrawal threshold (WT), measured in grams was multiplied by 10,000 and logarithmically transformed to obtain a linear scale. Tests were performed every other week by a blinded researcher.

### Measurement of anti-tAChR autoantibody levels

Antibody levels against tAChR in serum were determined using radioimmunoassay. Samples were diluted 1:100 (week 4) or 1:200 (week 8 and terminal timepoint) in PBS. A volume of 2.5 µL of diluted serum samples was incubated overnight with an excess of 0.5 µL ^125^I-α-bungarotoxin (BT) (Perkin Elmer, The US), 2.5 µL normal mouse serum and 197 µL tAChR ((0.25 µg/mL in PBS). Immune complexes were precipitated with 250 µL polyclonal goat-anti mouse Ig (Eurogentec, Belgium). The measured radioactivity of ^125^I-α-BT directly correlates to the amount of anti-tAChR antibodies present in serum and is calculated using the specific activity of the ^125^I-α-BT. All assays were performed in triplicate and the average of the measured radioactivity was used to calculate the antibody concentration expressed in nmol/L.

### Electromyography

Electromyography (EMG) measurements during curare infusion were recorded during terminal experiments. Animals received analgesia with 0.1 mg/kg buprenorphine and were anesthetized with isoflurane in the air. Body temperature was maintained at 35-37°C using an infrared lamp (DISA, Copenhagen, Denmark). Compound muscle action potential (CMAP) decrement was measured in the tibialis anterior (TA) muscle using the Viking IV EMG system (Nicolet Biomedicals, Madison, WI) as previously described ^31^. Briefly, the peroneal nerve was stimulated with a series of eight supramaximal stimuli of 3 Hz and a duration of 0.2 ms per stimulus. CMAPs were recorded from the TA. Decrement was confirmed when the fourth CMAP recording showed a ≥10% decrease in amplitude and area compared to the first CMAP in the same series, over three consecutive recordings. The amplitude primarily measures the strength of the response at a specific time, while the area captures the overall size and duration of the response. Measurements were repeated every 30 s, for three consecutive times as baseline measurements. In the absence of decrement, animals underwent tracheotomy to ensure proper ventilation before infusion of curare [(+)-tubocurarine, T2379; Sigma-Aldrich], acting as a competitive acetylcholine (ACh) antagonist. After making a small incision through the cutaneous and muscle layers of the peritoneum, a tube was then fixed in position (intraperitoneally; IP) using surgical tape to continuously infuse 20 μg/mL curare using a Terfusion syringe pump (model STC-521; Terumo, Tokyo, Japan) at a rate of 3.3 μL/min (0.066 μg curare/min). During curare infusion, CMAP measurements were repeated every 30 s until decrement was observed.

## Supplementary figures

**Supplementary figure 1.**
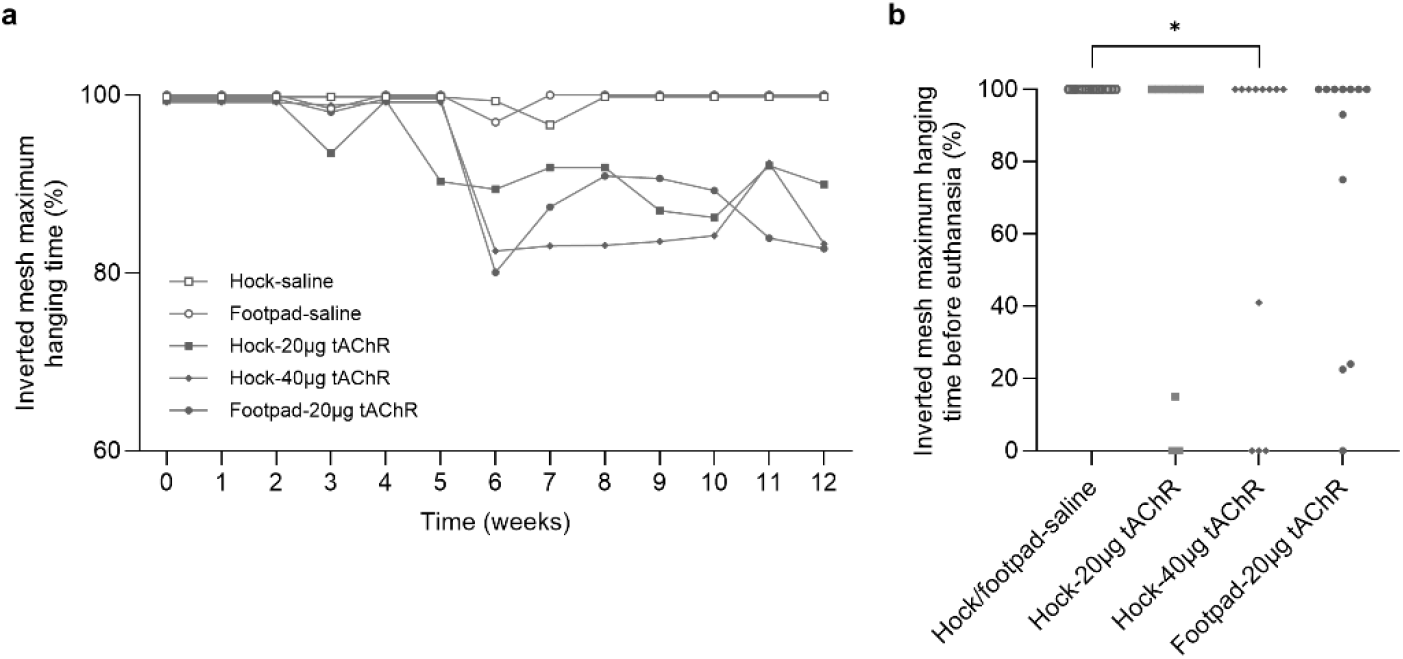
Inverted mesh hanging time. The maximum time spent by the animals in the inverted mesh (exclusively from the phase 1 experiment with reduced maximum hanging time of 320 s) as a % of the maximum hanging time for **a)** 12 consecutive weeks and **b)** individual hanging times at week 9. One and two-way ANOVA, multiple comparison of the indicated groups and Bonferroni post hoc testing were used for statistical analyses. *p<0.05.

**Supplementary figure 2.**
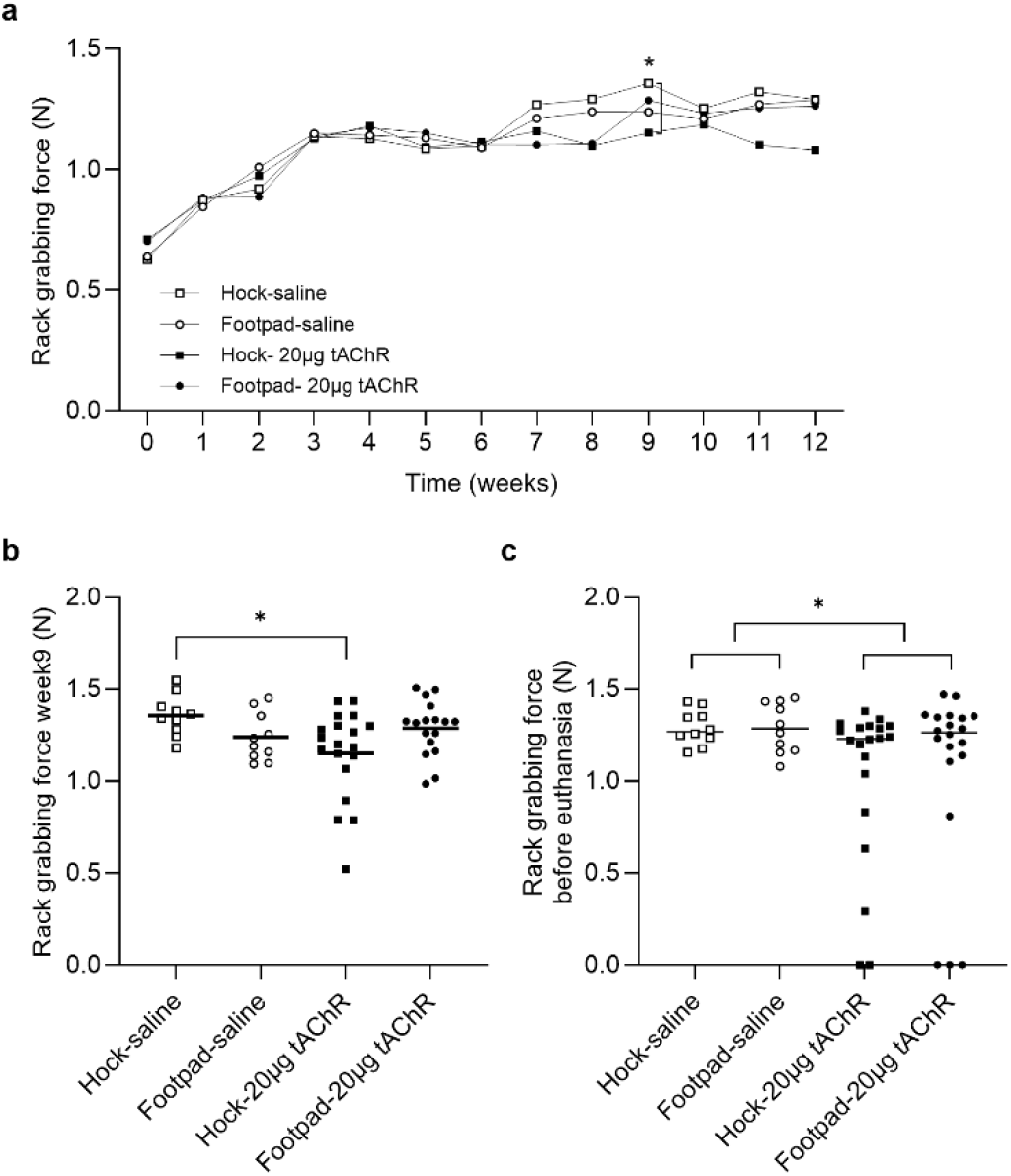
Rack grabbing force. **a)** Group-averaged rack grabbing force was measured weekly, **b)** individual values are shown at 9 weeks and **c)** at terminal timepoint. This analysis exclusively includes phase 2 experiment. One and two-way ANOVA, multiple comparison and Bonferroni post hoc testing were used for statistical analysis in A and B and Unpaired t test for C. *p<0.05.

## Acknowledgements

We thank Dr. Franken for his expertise and guidance on the Von Frey analysis.

